# Leptin activation of dorsal raphe neurons inhibits feeding behavior

**DOI:** 10.1101/2023.04.24.538086

**Authors:** N.D. Maxwell, C.E. Smiley, A.T. Sadek, F.Z. Loyo-Rosado, D.C. Giles, V.A. Macht, J.L. Woodruff, D.L. Taylor, S.P. Wilson, J.R. Fadel, L.P. Reagan, C.A. Grillo

## Abstract

Leptin is a homeostatic regulatory element that signals the presence of energy stores -in the form of adipocytes-which ultimately reduces food intake and increases energy expenditure. Similarly, serotonin (5-HT), a signaling molecule found in both the central and peripheral nervous systems, also regulates food intake. Here we use a combination of pharmacological manipulations, optogenetics, retrograde tracing, and *in situ* hybridization, combined with behavioral endpoints to physiologically and anatomically identify a novel leptin-mediated pathway between 5-HT neurons in the dorsal raphe nucleus (DRN) and hypothalamic arcuate nucleus (ARC) that controls food intake. In this study, we show that microinjecting leptin directly into the DRN reduces food intake in male Sprague-Dawley rats. This effect is mediated by leptin-receptor expressing neurons in the DRN as selective optogenetic activation of these neurons at either their ARC terminals or DRN cell bodies also reduces food intake. Anatomically, we identified a unique population of serotonergic raphe neurons expressing leptin receptors that send projections to the ARC. Finally, by utilizing *in vivo* microdialysis and high-performance liquid chromatography, we show that leptin administration to the DRN increases 5-HT efflux into the ARC. Overall, this study identifies a novel circuit for leptin-mediated control of food intake through a DRN-ARC pathway, utilizing 5-HT as a mechanism to control feeding behavior. Characterization of this new pathway creates opportunities for understanding how the brain controls eating behavior, as well as opens alternative routes for the treatment of eating disorders.

****Significance:**:** Leptin and serotonin both play a vital role in the regulation of food intake, yet there is still uncertainty in how these two molecules interact to control appetite. The purpose of this study is to further understand the anatomical and functional connections between leptin receptor expressing neurons in the dorsal raphe nucleus, the main source of serotonin, and the arcuate nucleus of the hypothalamus, and how serotonin plays a role in this pathway to reduce food intake. Insight gained from this study will contribute to a more thorough understanding of the networks that regulate food intake, and open alternative avenues for the development of treatments for obesity and eating disorders.

## Introduction

Leptin is a key homeostatic regulatory element that signals the presence of adipocyte energy stores and, under normal physiological conditions, reduces food intake and increases energy expenditure. To function in the brain, leptin crosses the blood brain barrier in a leptin-receptor-mediated fashion (1, 2), where leptin then binds to long-form receptors (LepRs). These receptors are expressed primarily on neurons within the hypothalamus (3), but also throughout the rest of the central nervous system (CNS) (4). Previously, mice with either knockout of the leptin gene (ob/ob) (5, 6) or knockout of functional leptin receptors (db/db) (7) elicits significant hyperphagia and severe obesity. LepRs are tyrosine receptor kinases that activate several pathways, most notably the JAK-STAT pathway (8). In particular, the phosphorylation of STAT3 is necessary for regulating energy homeostasis (9), by repressing the transcription of genes that increase hunger, such as agouti-related peptide (AGRP) and neuropeptide Y (NPY) (10, 11) while activating transcription of genes that suppress hunger, such as pro-opiomelanocortin (POMC) (12–14) and cocaine- and amphetamine-regulated transcript (CART). LepRs have been well characterized in the hypothalamus, the primary region of the brain that regulates homeostatic processes including appetite. Four hypothalamic nuclei have been identified as key regulators of food intake and energy expenditure: the arcuate nucleus (ARC), (15), lateral hypothalamus (LH), (16), the ventromedial nucleus (VMN), (17), and the paraventricular nucleus (PVN), (18). The anorectic effects of leptin through these neurons in the hypothalamus has been well described (19), however this is not the only route leptin can take to reduce appetite. For example, in addition to the hypothalamus, LepRs are expressed in several extra-hypothalamic areas, such as the ventrotegmental area (20) and the midbrain dorsal raphe nucleus (DRN) (3, 21, 22). However, the role of LepRs in the DRN has not yet been unequivocally determined.

The DRN is the main source of central serotonin (5-hydroxytryptamine,5-HT) (23). Functionally, 5-HT participates in the regulation of many behavioral and physiological processes through which energy balance is maintained (24). Regarding feeding behavior, 5-HT signaling plays a predominantly inhibitory effect and previous studies have shown that specific lesions or acute inhibition of raphe neurons cause hyperphagia and obesity (25, 26). In addition, inhibition of 5-HT synthesis by intracerebroventricular (icv) injection of either the serotonergic neurotoxin 5,7-dihydroxytryptamine (5,7-DHT) or the tryptophan hydroxylase (TPH) inhibitor p-chlorophenylalanine (PCPA) produces hyperphagia, further emphasizing that loss of 5-HT signaling exacerbates feeding behaviors (27, 28). Conversely, central injections of 5-HT or its precursor elicits hypophagia (29–32). Collectively, these findings support the critical role of 5-HT in attenuation of feeding behaviors.

The converging roles of leptin and 5-HT on food intake suggests a physiological integration, but the complexity of both systems throughout the brain convolutes the mechanisms by which they may regulate feeding behavior. For example, it has been postulated that 5-HT and leptin both represent independent energy-balance systems that are integrated in the hypothalamus, as, 5-HT type 2C receptors (5-HT2CRs) and LepRs are expressed in separate populations of POMC neurons in the ARC (33), suggesting that the integration of these systems may occur at another level. In support of this hypothesis, some studies suggest a complex interaction may occur between 5-HT and leptin systems within the DRN. Early histological studies showed that LepR are expressed in neurons of the raphe nuclei (3, 34, 35), where it co-localizes with SERT(36, 37). However, studies using an independently generated LepR reporter mouse model showed that raphe leptin receptors do not regulate food intake (38). Conversely, others have proposed a model in which leptin regulates appetite and energy expenditure through serotonergic circuits (39, 40). This model was based on transgenic mice in which LepR was specifically eliminated in SERT-expressing cells, producing a phenotype of increased food intake, body weight, and body fat along with decreased energy expenditure and bone mass. Collectively, these studies show discrepancies in the role of the LepRs in the raphe and the regulation of food intake. The aim of the current study was to discern the role of DRN LepRs on eating behavior and to establish the functional neuroanatomical connection between the DRN and the ARC of the hypothalamus. Although extensive studies have been done to define the projections from the raphe nuclei that reach different hypothalamic nuclei (41), including the ARC (42) there are no reports identifying raphe leptin-sensitive neurons that connect these nuclei. Using neuronal tract tracing, histology, pharmacological and optogenetics approaches, and *in vivo* microdialysis, we tested the hypothesis that leptin regulates food intake not only by activating hypothalamic LepRs, but also through activation of LepRs located in the serotonergic raphe neurons that send projections to the ARC.

## Results

### Section 1: Leptin reduces food intake through the DRN

To validate the efficacy of leptin utilized in our study, we injected leptin (or vehicle) both intraperitoneally (ip) and directly into the ARC, both of which have been shown to reduce food intake following leptin administration (5, 51, 52). We observed a reduction in food intake following a 5 mg/kg ip injection of leptin compared to vehicle, with a significant effect of leptin administration (*F*(_1,12_) = 68.06, *p* < 0.0001), significant effect of time (*F*(_5,60_) = 400.62, *p* < 0.0001), and a significant interaction on cumulative food intake (*F*(_5,60_) = 18.48, *p* < 0.0001) (Figure 1A). Further, post-hoc Bonferroni analyses showed significant differences in food intake between vehicle-treated rats and ip leptin-treated rats starting 3 hours post administration that persisted until at least 24 hours (3 hours: p = 0.0101; 4, 12 and 24 hours: p < 0.0001) (Figure 1A). Similar differences were observed when leptin was microinjected into the ARC, with a significant effect of leptin administration (*F*(_1,8_) = 50.67 *p* < 0.0001), significant effect of time (*F*(_6,48_) = 114.96, *p* < 0.0001), and a significant interaction on cumulative food intake (*F*(_6,48_) = 11.10, *p* < 0.0001) (Figure 1B). Therefore, acute administration of leptin caused a delayed and long-lasting inhibition of food intake both peripherally and intra-ARC. To determine if DRN leptin controls food intake, we injected varying concentrations of leptin directly into the DRN. We observed a dose-dependent reduction of food intake. Notably, following an injection of 1 µg of leptin directly into the DRN, we observed a reduction in food intake equivalent to that observed in a peripheral or icv injections (Figure 1C). Two-way repeated measures ANOVA on cumulative food intake revealed significant main effects of intra-raphe leptin administration (*F*(_31,68_) = 5.58, *p* = 0.0082), a significant effect of time (*F*(_7,12_) = 125.21, *p* < 0.0001), and a significant interaction (*F*(_21,112_) = 5.82, *p* < 0.0001) on the cumulative food intake (Figure 1C). Post-hoc Bonferroni analyses revealed that 1 µg of leptin significantly inhibited food intake 4 hours after hormone infusion and this effect was sustained for at least 24 hours, whereas administration of lower doses failed to decrease food intake.

**Figure 1.**
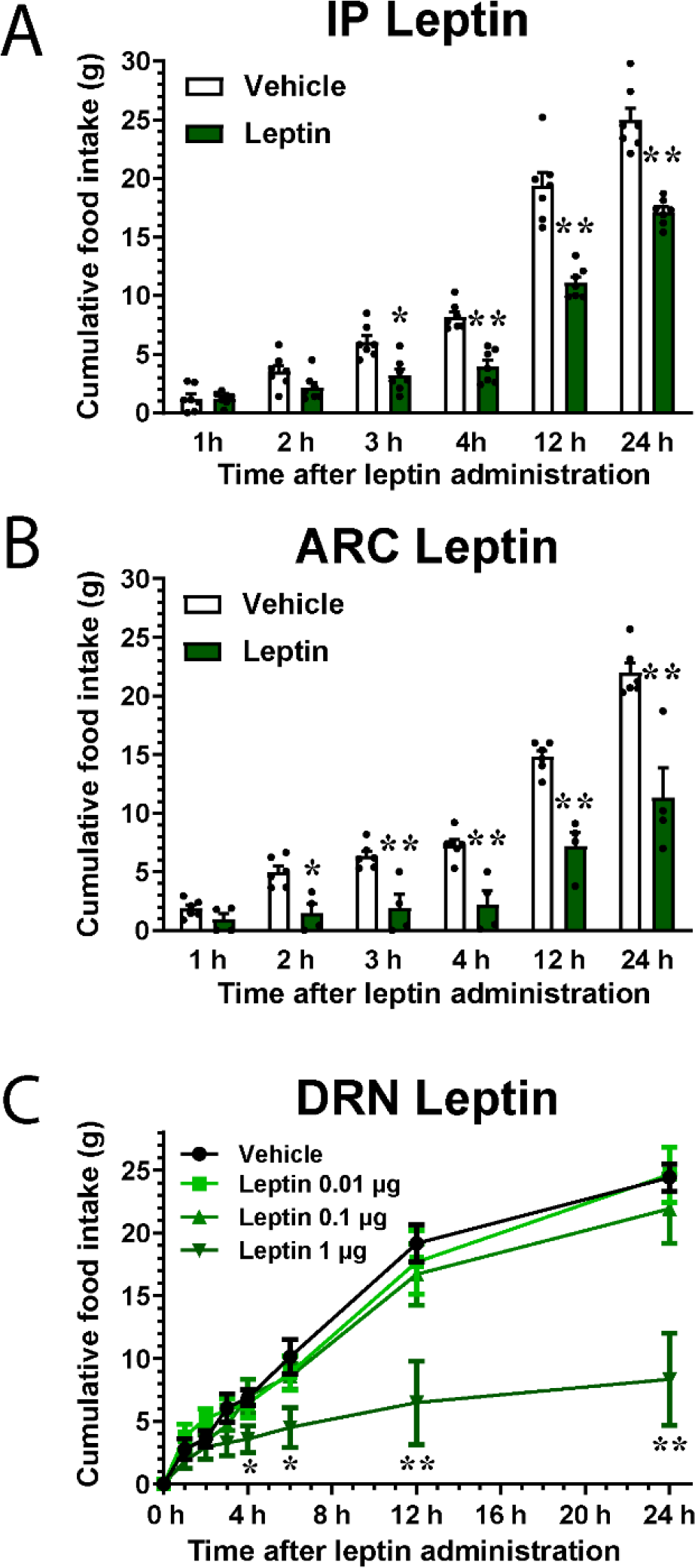
Leptin inhibits food intake through peripheral, arcuate (ARC), and dorsal raphe (DRN) injections. (A) 5 mg/kg leptin was ip injected at the beginning of the dark cycle, and hoppers were weighed at 1, 2, 3, 4, 12 and 24 hours post injection (n=7 for each group). (B) 1 µg of leptin (n=4) or vehicle (n=6) was injected into the ARC, and food hoppers were weighed as before. (C) Microinjections of vehicle, 0.01 µg, 0.1 µg, and 1 µg were delivered to the DRN (n=5 for each group). Error bars = SEM. Significance at specific time points were identified using a post-hoc Bonferroni’s multiple comparisons test. *p <0.05, **p<0.01 compared to vehicle treated animals.

### Section 2: Leptin sensitive DRN neurons project to the ARC

Following validation of leptin’s ability to control food intake both intra-ARC and intra-DRN, we explored the connection between these two regions. Using injections of Alexafluor conjugated neuronal retrograde tracer, cholera toxin subunit b (fCTb), we show an anatomical connection between the DRN and the ARC (Figure 2A). Following injection of fCTb into the ARC, the tracer is taken up by axonal terminals in the ARC and retrogradely transported where it accumulates in the cell bodies of projecting neurons, including projecting neurons in the DRN (Figure 2A-C). From rats that were given 5 µg leptin icv, DRN sections were then immunolabeled for phosphorylated STAT3 (pSTAT3), a commonly used marker of leptin signaling. Some neurons in the DRN that express pSTAT3 are also positive for fCTb (Figure 2D), which establishes the presence of DRN leptin receptor expressing neurons that project to the ARC, potentially to control appetite.

**Figure 2.**
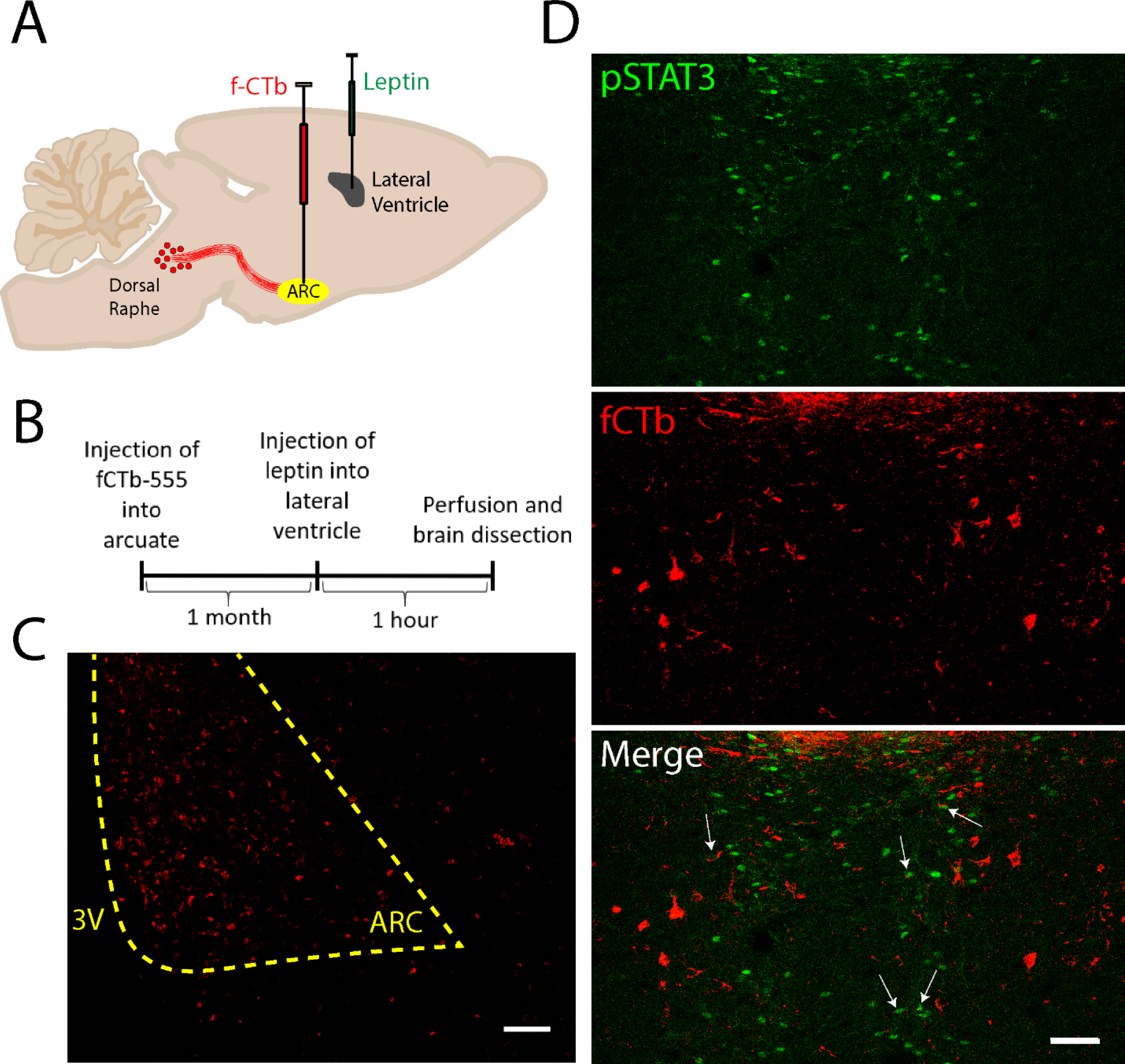
Retrograde tracing shows that leptin sensitive DRN neurons project to the ARC. (A,B) Created with BioRender.com. Experimental design and timeline for the experiment. Fluorescently labeled CTb-555 is injected into the ARC and left for 1 month, then leptin is injected 1 hour before sacrifice to induce STAT3 phosphorylation. (C) The injection site of CTb-555 is shown in the ARC at 10×. (D) 20× confocal image of the DRN, with leptin sensitive neurons (pSTAT3, green) and retrograde-labeled neurons that project to the ARC (fCTb, red). White arrows note co-labeled neurons, indicating activation of LepR on ARC projecting neurons. Scalebars = 50µm.

### Section 3: Optogenetic stimulation of leptin receptor expressing neurons in the DRN reduces food intake

To establish a functional role of the LepR expressing neurons in the DRN, we utilized an optogenetic approach to target LepR expressing neurons in the DRN. We designed a lentivirus that would exclusively express the light-sensitive protein channelrhodopsin-2 (ChR2), a transmembrane ion channel activated by 470nm light, in neurons that express the long form of the leptin receptor (LV-ChR2), as well as a control virus lacking the ChR2 sequence (LV-Con, Figure 3A). In animals injected with LV-Con into the DRN, there was no alteration in food intake compared with previous food intake trials, nor was there any changes in food intake based on the wavelength of light given (*F*(_12,66_) = 0.92, p = 0.5303) (Figure 3B). However, when LV-ChR2 was injected into the ARC, blue light-ARC significantly reduced the amount of food consumed compared to yellow light-ARC (*F*(_3,72_) = 22.36, p < 0.0001) (Figure 3C). Then, to explore the role of LepR expressing neurons in the DRN, LV-ChR2 was injected into the DRN and blue and yellow light administered to the DRN. Again, food intake was significantly decreased in the rats that received blue light-DRN stimulation as compared to yellow light-DRN stimulation over time (*F*(_6,60_) = 418.49, p < 0.0001) and cumulatively (*F*(_6,10_) = 5.10, p < 0.0003) (Figure 3D). These results support the hypothesis that LepR-expressing neurons in the DRN reduce food intake. To further determine whether these reductions in food intake occur through projections to ARC neurons, lentivirus was injected into the DRN, but the optogenetic probe was placed into the ARC. Photostimulation of the axon terminals in the ARC with blue light significantly reduced food intake compared to yellow light-ARC administration over time (*F*(_6,48_) = 602.94, p < 0.0001) and cumulatively (*F*(_6,48_) = 2.89, p = 0.0173) (Figure 3E), similar to was observed following direct stimulation of DRN somata.

**Figure 3.**
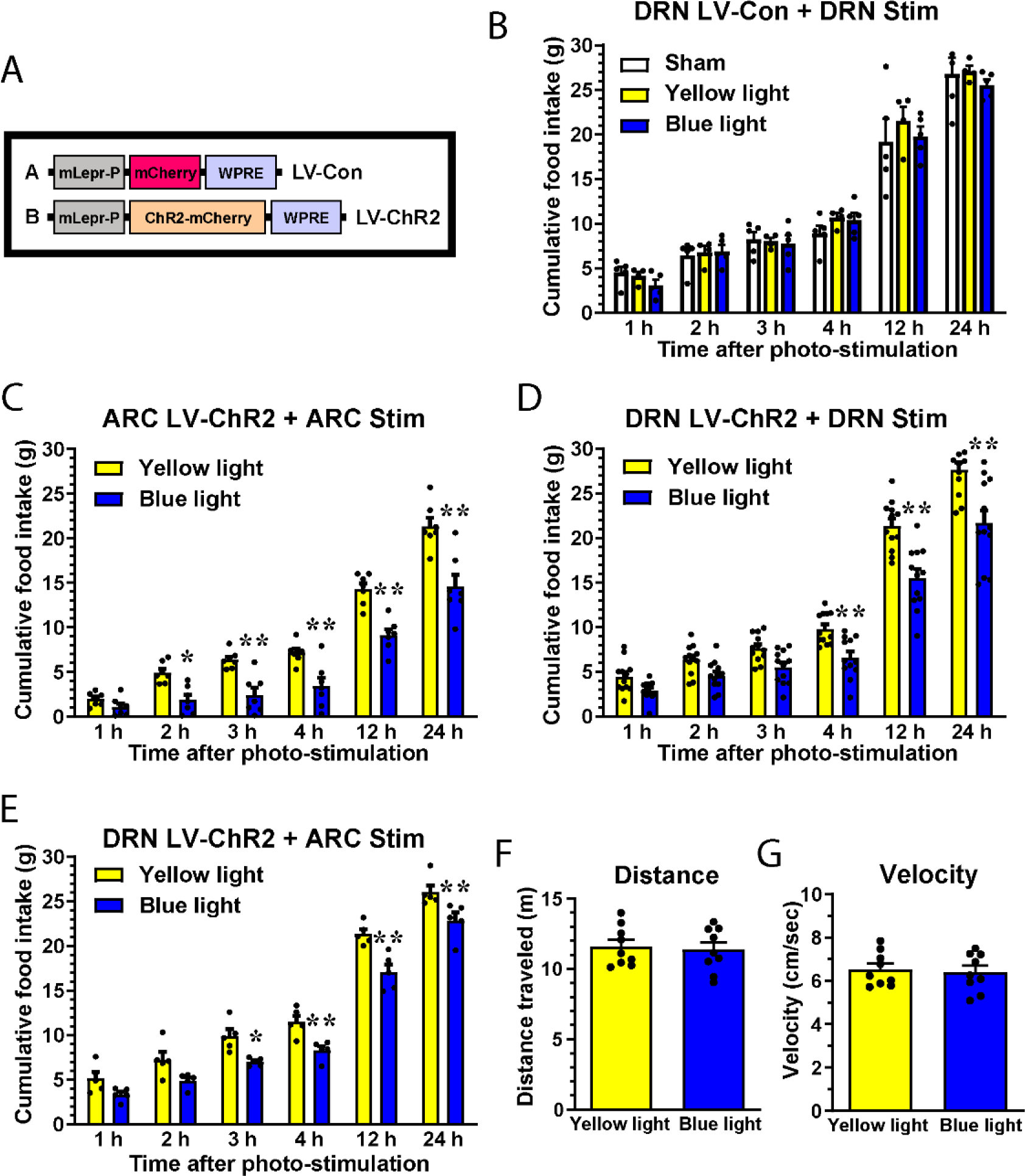
Optogenetic stimulation of leptin sensitive neurons in the DRN induces a reduction in food intake. (A) Lentivirus was used to express channelrhodopsin-2 (ChR2) in LepR expressing neurons. Control virus only expresses mCherry under the LepR promoter, the experimental virus expresses a ChR2 mCherry fusion protein under control of the LepR promoter. (B) Food intake recording following stimulation of the DRN infected with LV-Con with blue light (470nm) (n=5), yellow light (590nm) (n=4), or no light (sham, n=5). (C) Food intake recording following stimulation with blue light (or yellow light as a negative control) of the ARC in animals with LV-ChR2 injected into the ARC (n=7 for both groups), (D) stimulation of the DRN with LV-ChR2 injected into the DRN (n=12 for both groups), or (E-G) stimulation of the ARC with LV-ChR2 injected into the DRN (n=5 for both groups). (F, G) Measurement of ambulatory function in an open field following photostimulation of leptin receptor expressing neurons of the DRN with yellow or blue light (n=9 for both groups). *p <0.05, **p<0.01 compared to yellow light treated animals.

To determine if activation of these leptin-sensitive neurons of the DRN affects locomotor activity, we subjected rats that were injected with LV-ChR2 into the DRN to open field test after photostimulation with blue and yellow light. No changes in locomotor activity with regards to distance traveled (p = 0.7219) or velocity of travel (p = 0.7271) were observed (Figure 3 F-G) suggesting that any effects of photostimulation on food intake are not due to changes in mobility.

### Section 4: Leptin-responsive serotonergic DRN neurons send projections to the ARC

To further explore the potential relationship between 5-HT and leptin in controlling food intake, we injected a fluorescently conjugated retrograde tracer (fCTb) into the ARC and combined with immunofluorescent markers for activation of LepR signaling (i.e., pSTAT3) and serotonergic neurons (TPH). We observed that serotonergic neurons from the DRN project to the ARC, and that leptin activates a subset of these projecting neurons (Figure 4A). Then, using LV-Con to label LepR expressing DRN neurons with mCherry, projections to the ARC were observed adjacent to immunostaining for serotonin reuptake transporter (SERT), which is located in serotonergic synaptic regions (Figure 4B). Colocalization between mCherry labeled projections and SERT labeling in the ARC are indicative of projections from serotonergic LepR expressing DRN neurons (Figure 4C).

**Figure 4.**
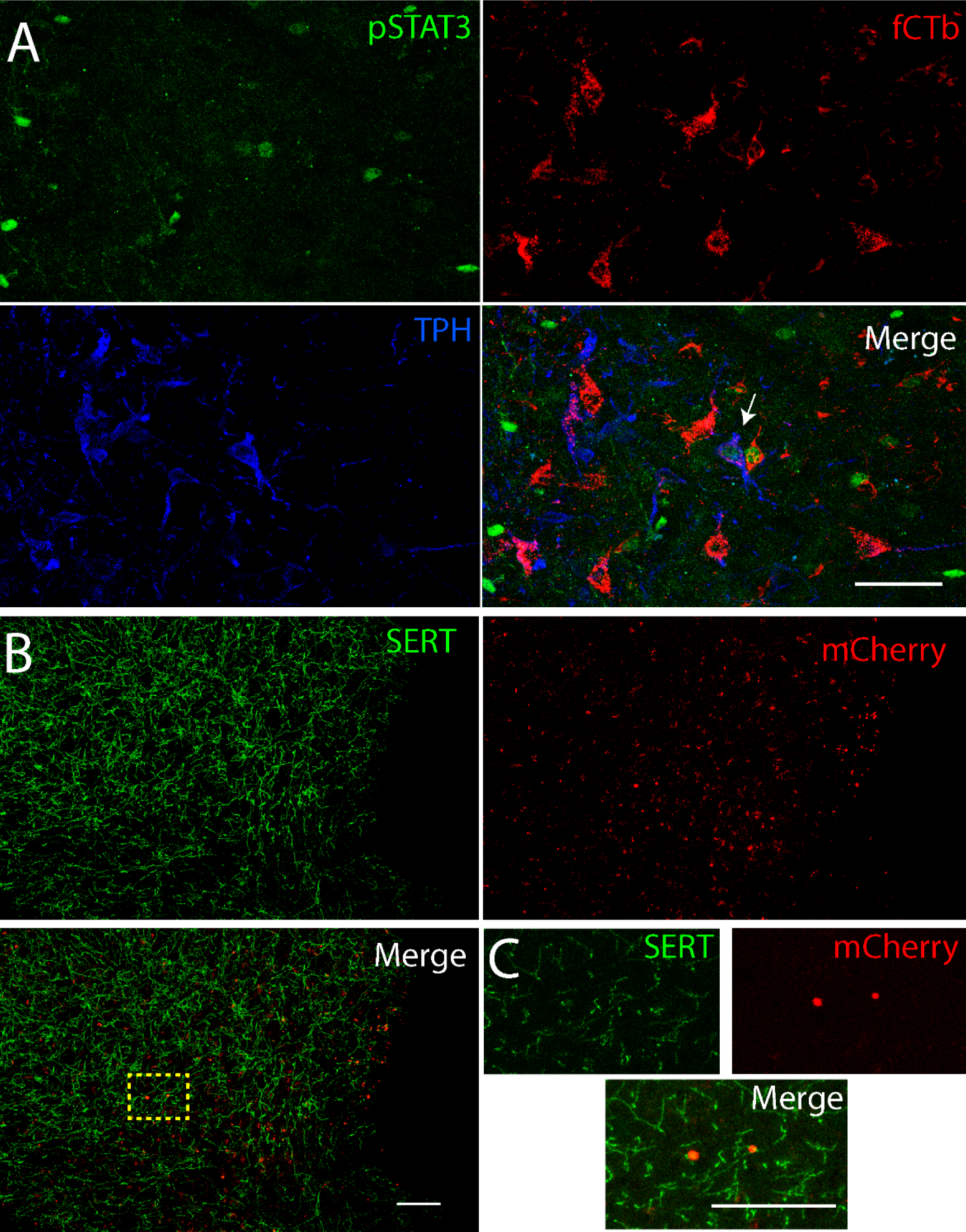
Serotonergic neurons sensitive to leptin project from the DRN to the ARC. (A) Fluorescently labeled CTb-555 was injected into the ARC and left for 1 month, then leptin was injected 1 hour before sacrifice to induce phosphorylation of STAT3 in the DRN. Confocal image taken with a 20× objective of the DRN, with leptin sensitive neurons (pSTAT3, green), neurons that project to the ARC (fCTb, red), and serotonergic neurons (TPH, blue). White arrow notes triple labeled neurons. (B) Image of the ARC taken with a 10× objective, where there are projections from leptin expressing neurons in the DRN (mCherry, red), and serotonergic reuptake transporter (SERT, green). (C) Image of ARC taken with a 40× objective showing colocalization of projections from leptin receptor expressing DRN neurons and SERT. Scalebars = 50µm across all images.

### Section 5: LepR-expressing serotonergic DRN neurons send axons to the ARC

We utilized RNAScope (fluorescent in situ hybridization) to label neurons that express LepR mRNA. Combining this technique with our previously described retrograde tracing and immunofluorescence, we were able to confirm the strong presence of LepRs in the DRN. We also identified multiple serotonergic neurons that project to the ARC and contain LepR mRNA (Figure 5A-B). Thus, we identified leptin sensitive neurons using two different approaches: 1) by expression of p-STAT3 (marker for LepR activation) and 2) expression of the mRNA for the leptin receptor. Moreover, in both cases, these LepR-expressing neurons exhibited projections to the ARC.

**Figure 5.**
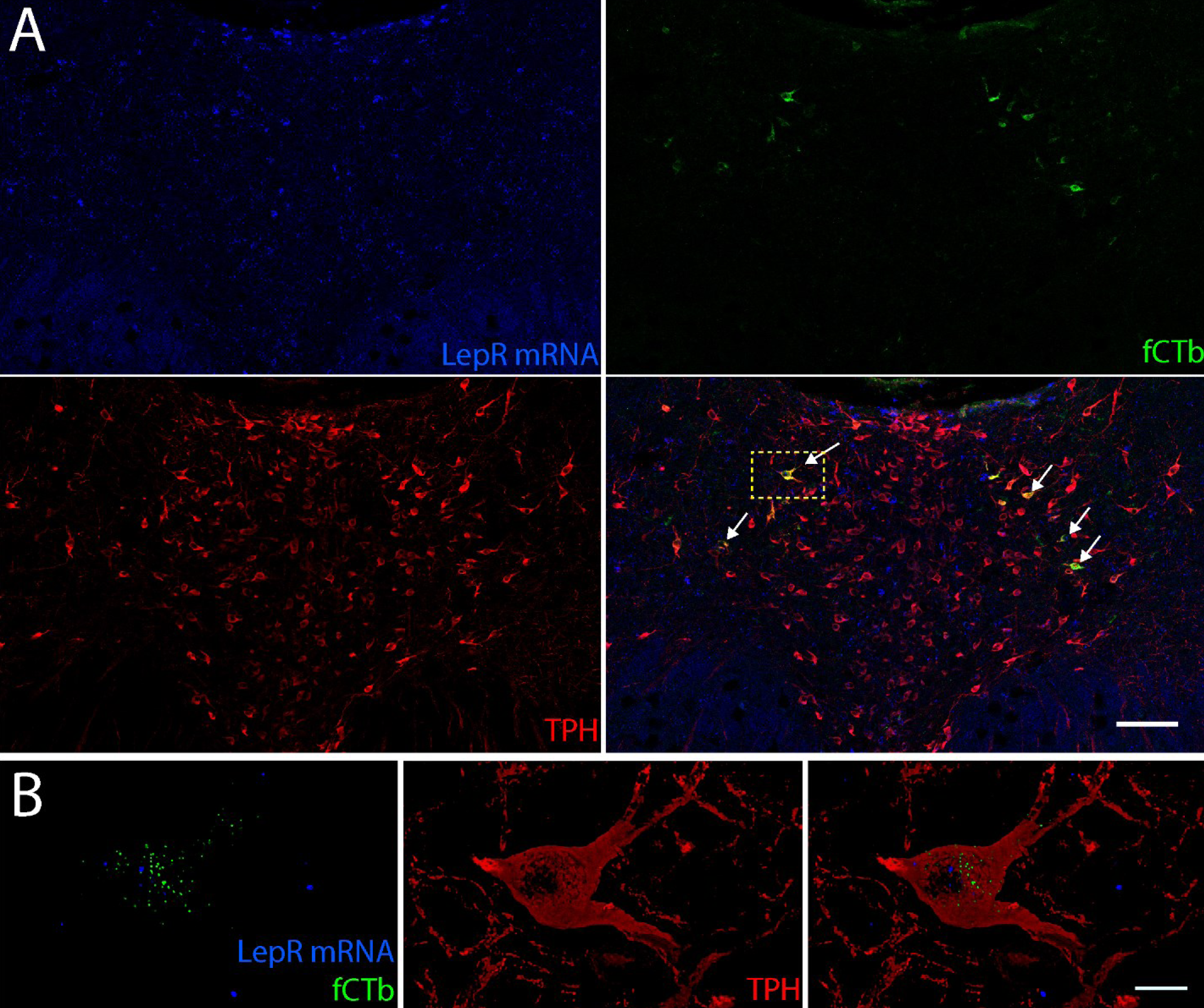
Leptin receptor mRNA is expressed in serotonergic DRN neurons that project to the ARC. (A) A 20× tile scan image of the DRN. Leptin receptor mRNA is visualized using RNAScope and opal 650 dye (blue). Neurons that project to the ARC are visualized with fCTb (green), and serotonergic neurons are visualized with immunofluorescence against TPH (red). Neurons showing LepR mRNA, fCTb, and TPH are noted with white arrows. (B) 63× Lightning super-resolution image of a serotonergic neuron with LepR mRNA (blue), fCTb (green), and TPH (Red). Scalebars = 500µm with 20× objective (A) and 5µm with 63× objective (B).

### Section 6: Activation of DRN LepRs stimulates the efflux of 5-HT in the ARC

To test our hypothesis that leptin stimulation of serotonergic raphe neuros drives release of 5-HT in hypothalamic nuclei, we microinjected leptin into the DRN and collected microdialysates in the ARC. Administration of leptin to the DRN showed a statistically significant increase of 5-HT efflux in dialysate collected from the ARC compared to vehicle administration, specifically at collection 8 (p = 0.0476) and 13 (p = 0.0072) using a Bonferroni’s multiple comparisons test. Further, there was a significant effect of time x treatment interaction (*F*(_19, 190_) = 1.96; DFn = 19; DFd = 190) and treatment (*F*(_1, 10_) = 6.70; DFn = 1; DFd = 10) (Figure 6B). The analysis of the area under the curve of the 5-HT efflux also showed statistically significant increases after leptin administration compared to vehicle treatment (p = 0.0278, Figure 6C-D).

**Figure 6:**
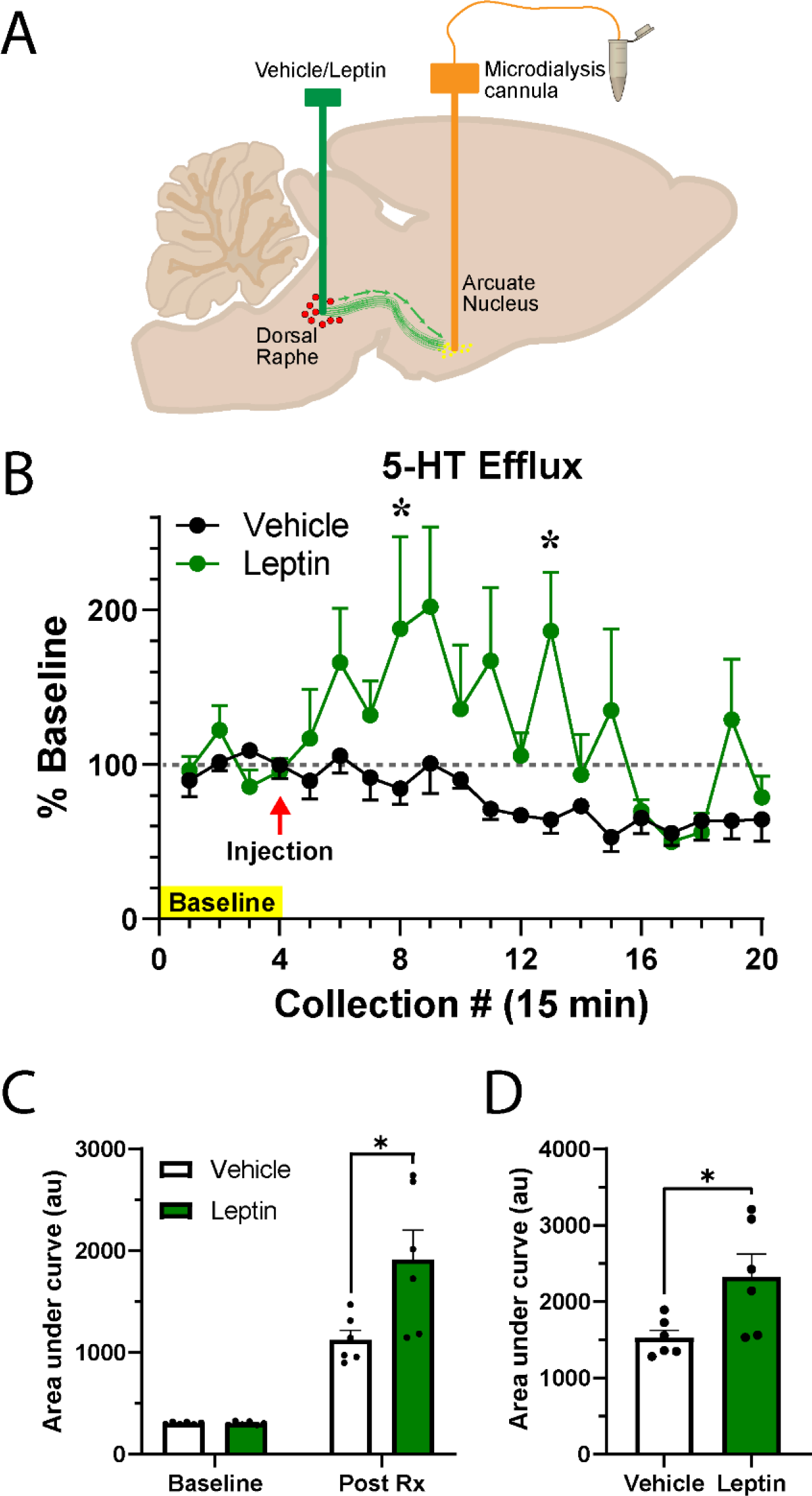
Administration of leptin to the DRN increases ARC 5HT efflux. (A) Diagram of cannula placement within the DRN for leptin infusion, and the ARC for the dialysis probe. (B) Changes in 5-HT efflux from baseline following vehicle (black) or leptin (green) injection at the end of collection 4. (C) Area under the curve for 5-HT efflux shown during baseline and after treatment, (D) as well as total area under the curve between vehicle and leptin treated animals. n=6 for each group. *p <0.05.

## Discussion

The current study used converging pharmacological, physiological, and anatomical techniques to identify the mechanisms by which DRN neurons that express LepR mediate feeding behavior. The results of the current study demonstrate that the DRN plays a role in feeding behavior in rats. Specifically, our data show that intra-DRN administration of leptin effectively suppresses food intake. In addition to the exogenous administration of leptin, optogenetic activation of the leptin-sensitive neurons located in the DRN also inhibits food intake and this effect was also evoked when the axonal terminals that reach the ARC were opto-stimulated. Collectively, these results support the hypothesis that leptin suppresses food intake by acting upon the DRN, and in turn the DRN is capable of releasing 5-HT into the ARC and regulating feeding behavior through the leptin-sensitive serotonergic projections from the DRN to the ARC. These data are the first to show that the LepR-expressing neuronal population within the DRN is sufficiently able to reduce food intake without direct stimulation of hypothalamic neurons by leptin or 5-HT. Moreover, a subset of this neuronal population is serotonergic supporting a novel link between leptin and 5-HT to suppress food intake.

Despite the wide distribution of LepRs in the rat brain (3), the study of the central control of food intake by leptin has been primarily focused on the leptin receptors located in the hypothalamus (17). However, cell-specific deletion of LepRs in the ARC of the hypothalamus (15, 17) does not prevent the anorexigenic effect of leptin in mice. In addition, deletion of leptin-sensitive neurons in the ventromedial nucleus of the hypothalamus of rats attenuates the food intake inhibition but does not eliminate it (53), suggesting that the food intake control by leptin is not confined only to the hypothalamus. In support of this, stimulation of hindbrain leptin receptors decreases food intake and body weight in rats (54), and a more recent study also in rats showed that decreasing the leptin receptor-expressing neurons in the hindbrain attenuates the response to leptin (55), supporting the notion that leptin regulates energy balance through a circuit beyond the hypothalamus, more precisely the hindbrain. Using a LepR-IRES-Cre EYFP reporter mouse, Scott et al. (2009)(22) found substantial hypothalamic and extrahypothalamic expression of the LepR. Among the latter, the DRN shows a high density for the mRNA and robust YFP expression, and an important co-localization with pSTAT3 (22). Concomitantly, using a transgenic mouse co-expressing LepR and enhanced green fluorescent protein (EGFP), Patterson et al. (2011) (56) found abundant expression of LepR in the soma of neurons located in the forebrain, midbrain, and hindbrain. As expected, the hypothalamic nuclei showed the highest expression; nonetheless, among the midbrain nuclei the DRN showed a substantial expression. Undoubtedly, while the DRN is a target for leptin, the physiological role of leptin in this nucleus is still unclear. Although a few studies suggest the regulation of food intake by leptin action in raphe neurons, the mechanisms underlying this phenomena remain unclear (57). Therefore, we focused our attention on the DRN, an area of the brainstem that was previously explored but with disparate results.

One of the pioneer studies to test the effects of leptin administered directly into the raphe nuclei on food intake and body weight in rats showed that small doses of leptin were able to suppress food intake when injected into the hypothalamic ventromedial nuclei, but this effect did not occur when leptin was injected into the lateral ventricle or the DRN (58). It is noteworthy that in that study, 0.05 µg leptin was infused twice daily; in our laboratory with a similar dose (0.1 µg) no significant effects were observed, but rather an injection of 1 µg of leptin was necessary to obtain a significant reduction in feeding behavior. These results suggest that even though leptin receptors are widely spread in the CNS, it is possible that different nuclei have varying levels of sensitivity to the hormone, and region-specific activation depends on leptin levels. When leptin was administered directly into the 4^th^ ventricle, the suppression of food intake was significant with a dose of 0.83 µg, whereas 0.1 µg of leptin was not sufficient to regulate feeding behavior or body weight changes in rats (59). Subsequent studies show that chronic infusions of subthreshold doses of leptin into the 3^rd^ and 4^th^ ventricles are able to decrease food intake and body weight when they are applied simultaneously in both ventricles and there is no effect when rats received leptin into either the 3^rd^ or the 4^th^ ventricle (60). These results suggest that hindbrain LepRs play a crucial role in the regulation of feeding behavior, particularly at higher levels of leptin concentration. Despite all these observations, few studies to elucidate the specific role of the DRN LepR upon food intake have appeared. Yadav et al. (2011) (40), using a mouse model lacking LepR in serotonergic neurons, found that the mice developed hyperphagia and obesity. Whereas Lam et al. (2011) (38), found that leptin’s ability to produce hypophagia does not depend on 5-HT bioavailability in mice.

To our knowledge this is the first study to determine the role of the rat DRN on feeding behavior. The administration of leptin directly into the DRN dose-dependently suppressed food intake. In addition, the leptin effect in the DRN exhibits a similar pattern compared with the regulation of appetite observed when the hormone is administered directly into the ARC or peripherally. Regardless of the route of administration, leptin causes a delayed and long-lasting effect that was reproduced using an optogenetic approach. For this purpose, we designed a lentivirus that encodes the light-sensitive ion channel protein channelrhodopsin-2 (ChR2) coupled to mCherry (reporter gene) under the LepR promoter. This construct allowed us to selectively target the leptin-responsive neurons in the DRN. This approach has some limitations as expression of ChR2, and thus activation of the neurons, is dependent on the viral spread through the tissue. Second, neuron activation is limited by the penetrance of the administered light. These limitations actually suggests that we are underestimating the anorexigenic effect of leptin compared to the microinjections of leptin. The results from both pharmacological and optogenetic approaches are congruent: activation of DRN LepRs suppresses food intake. At the same time, the optogenetics approach allowed us to identify the projections from the DRN that participate in the anorectic effect of leptin. In the current study, we have shown that DRN projections sent to the ARC are certainly involved in this circuit; however, DRN projections to other hypothalamic nuclei, such as LH or VMN, could be as vital as this DRN-Arc pathway, and is currently being investigated in our laboratory.

Given that 5-HT is the primary neurotransmitter synthesized in the DRN (23), we then evaluated if 5-HT is a driver of changes in food intake following intra-DRN leptin administration. Serotonin regulates many behavioral and physiological processes through which energy balance is maintained, playing a predominantly inhibitory effect regarding feeding behavior (24). Lesions or inhibition of raphe neurons resulted in hyperphagia and obesity (25, 26) and inhibition of 5-HT synthesis by icv injection of either the serotonergic neurotoxin 5,7-dihydroxytryptamine (5,7-DHT) or p-Chlorophenylalanine (PCPA) also produces hyperphagia in rats (27,28). Conversely, central injections of 5-HT or its precursor elicits hypophagia (29–32). In C57/BL6 mice, the deletion of TPH2 promotes hyperphagia and body weight gain in males, but not in females (61). In another report investigating the interaction of 5-HT and leptin to influence food intake, the depletion of 5-HT did not affect the leptin inhibitory effect on food consumption (38). It is important to note that the PCPA and leptin treatments in this last study were done peripherally while our treatments were done directly into the DRN.

In the current study, we targeted a sub-population of raphe neurons that are activated by leptin and observed that some of them also produce 5-HT, supporting the notion that 5-HT is likely a major contributor to this pathway. Contrary to Lam et al. (2011) (38) but in accordance with Fernandez-Galaz et al. (2002) (36), we observed co-localization of TPH and pSTAT3 after leptin activation. Additionally, we confirmed this co-localization using RNAScope to label mRNA for the leptin receptor in DRN neurons. Furthermore, projections to the ARC from LepR expressing neurons in the DRN localize to SERT, the protein responsible for termination of 5-HT signaling. Although the neuroanatomical connections between the DRN and the hypothalamus have been described (42, 63), here we expand upon these initial observations to demonstrate functional connections between the serotonergic raphe neurons that project to the ARC and a novel circuit by which leptin controls feeding behavior.

Within this study, we show that administration of leptin to the DRN induces 5-HT release into the ARC. By microinjecting leptin specifically into the DRN and utilizing microdialysis to collect 5-HT from the ARC, all leptin-receptors expressing neurons within the DRN that project to the ARC are evaluated. This is in contrast to previous studies that have looked at the firing patterns of individual neurons within the DRN in response to leptin, thus potentially omitting critical groups of neurons in this pathway (38). This data definitively shows that leptin stimulates the 5-HT system in controlling the ARC. In the current study, we explored only serotonergic neurons in the DRN as targets of leptin activity, as they are the most abundant neuronal phenotype in that region (23). However, other neurotransmitters may play a valuable role in leptin’s ability to regulate food intake through the DRN and remain an important area for future investigation.

In conclusion, our studies show that there is an anatomical and functional connection between the DRN and the ARC of the hypothalamus, and that this connection is, at least in part, responsible for leptin’s ability to regulate food intake in rats. The characterization of this pathway firmly shows that leptin works outside of the hypothalamus to control food intake and identifies a new level of interaction between leptin and serotonin to regulate food intake. This also exposes alternate pathways that may be exploited in the future to develop treatments for eating disorders.

## Materials and Methods

### Animals

Adult male Sprague Dawley rats (CD Strain, Envigo), 2 months of age, weighing around 200-250 g were housed individually for the duration of the experiments. Rats had ad libitum access to food (standard chow) and water and were housed in a reverse 12-hour light/dark cycle (lights off at 11:00AM). All procedures involving animals were carried out in accordance with guidelines and regulations of the University of South Carolina Animal Care and Use Committee.

### Leptin treatment

Recombinant mouse leptin from the National Hormone and Peptide Program (NIH-NIDDK) was administered to the rats as a peripheral injection (5 mg/kg of body weight (BW)), icv infusion (5 µg), or intra-ARC administration (1 µg), consistent with previous literature (43). For intra-raphe administration each rat received 0.01, or 0.10, or 1 µg of leptin. Sterile 0.1 M phosphate buffered saline (PBS) was used as vehicle for all injections.

### Food intake measurements

Animals were habituated to food hoppers for a minimum of three days prior to experiments and had ad libitum access to food and water. Prior to surgeries, baseline food intake measurement began at the start of the dark cycle at 1, 2, 3, 4, 6, 12, and 24 hours. All experimental food intake measurements were taken at the same time intervals.

### Stereotaxic surgical coordinates

Rats were handled daily for 5 days prior to stereotaxic surgery. All injection coordinates were determined using the Paxinos and Watson brain atlas (1998). Rats were anesthetized with isoflurane (5% induction and 2-3% for maintenance) and placed in the stereotaxic apparatus (David Kopf Instruments). For leptin or lentivirus injections and optogenetic probe placement, the ARC was targeted using the following coordinates: AP: -2.6 mm; L: ±0.3 mm; DV: -10.0 mm. To target the dorsal raphe, the following coordinates were used: AP: -7.8 mm; L: 0.0 mm; DV: -6.0 mm. For injection cannula placement, the same coordinates were used, but 1mm more dorsal to account for the length of the infusion injector.

### Lentivirus production

A lentivirus transfer vector designed to express a channelrhodopsin (hChR2)-mCherry fusion protein under control of the mouse leptin receptor promoter was constructed using standard molecular cloning techniques. Briefly, a DNA fragment containing 863-bp of the mouse leptin receptor promoter (44) was amplified from a plasmid generously provided by Dr. Tetsurou Satoh (Gunma University, Maebashi, Japan). This promoter-containing fragment was then cloned into a promoter-less lentivirus transfer vector based on a third generation, self-inactivating transfer vector (45). The hChR2-mCherry fusion protein cDNA (46) was subsequently cloned into this transfer vector. This lentivirus was designated LV-ChR2. A control virus, LV-Con, was similarly constructed with the leptin receptor promoter controlling expression of mCherry (Clontech) without ChR2. Viruses were generated by transfection of the transfer vector with three packaging plasmids (Addgene), psPAX2, pRSV-Rev and pMD2.G, into 293T cells. Viruses were concentrated by high-speed centrifugation, purified by further centrifugation through 20% sucrose/PBS-D, and stored in 10% sucrose/PBS-D at -80°. Virus particle concentrations were determined by quantitative real-time PCR for proviral DNA 24 hours following transduction of 293T cells and are expressed as transducing units per microliter (tu/µl). A minimum of three weeks was given to allow for viral expression after injection.

### Stereotaxic surgeries for lentivirus injection and fiber optic implant

Using the same procedure described previously to target the ARC and the DRN, rats were injected with a lentivirus that encodes the light-sensitive ion channel ChR2-mCherry fusion protein under control of the LepR promoter (LV-ChR2). Control rats received the control construct (LV-Con) that encodes on mCherry but not ChR2. Five µL of the viral stock (5×10^6^ tu/µL) were injected at a speed of 1.0 μL/min for 5 minutes with a 10 μl Hamilton syringe driven by a motorized stereotaxic injector (Stoelting 53310); the needle was left in place for additional 15 min post-injection. Three weeks after the lentivirus infusion, the rats were subjected to a second stereotaxic surgery to implant the optical fiber cannula (400 µm core diameter, 0.39 NA, length: 10 mm, Thorlabs, Newton, NJ) through the same craniotomy as the viral injection using the same coordinates. Animals were allowed 3 days post-surgery to recover before behavioral assessments.

### Optical activation

Implanted fiber optic probes were connected to fiber optic cables leading to the stimulating light source (470nm wavelength, Thorlabs, M470F1) or the control light source (590nm wavelength, Thorlabs M590F1) and LED emitters (Thorlabs DC2100). Rats were then stimulated in their home cages for 5 minutes (150 Amp current, 3 Hz frequency (square wave), 50% duty cycle). Following stimulation, fiber optic cable leads were disconnected, and food was placed into the cage at the beginning of the dark cycle and weighed as described above.

### Open field test

Rats were subjected to an open field test to evaluate general locomotor activity following LV-ChR2 activation in the DRN. The open field apparatus consisted of a square, black plexiglass box (76×76×46 cm) with the floor divided into 25 squares (15×15 cm). The box was electronically divided into an inner square and an outer area surrounding the inner square. Animals were placed into the center of the open field 1 hour after photo-stimulation with yellow or blue light into the DRN and allowed to explore freely for 30 min. Data were collected using the Ethovision (Noldus, Leesburg, VA) automated system. The box was cleaned with alcohol before each testing session to minimize the scent of other rats. Measurements included total distance, and velocity.

### Retrograde tracing and immunofluorescence

For each study, injection sites were confirmed using fluorescent confocal microscopy (for retrograde tracer). Rats were anesthetized using vaporized isoflurane and bilaterally injected with 5 µg of Alexafluor-555 labeled fluorescent choleratoxin subunit b (fCTb) (Invitrogen C34776) into the left and right side of the ARC using the following coordinates from bregma: A/P: -3.14 mm; L: ±1.86 mm; D/V: -10.15 mm at an angle of 9° away from the midline. fCTB was injected at a concentration of 10 µg/µL at a speed of 0.1 µL/min using a 1 µL syringe driven by motorized stereotaxic injector (Stoelting 53310). The needle was left at the target for an additional 10 minutes to allow for settling of injected toxin before removal. Total volume injected into each side was 0.5 µL. Three weeks later, the rats were stereotaxically injected with 5 µg of leptin into the lateral ventricle: A/P: -0.8 mm; L: -1.4 mm; D/V: -3.8 mm at a speed of 1 µL/min over 5 minutes. One hour after the leptin infusion, the rats were transcardially perfused with 0.1 M phosphate buffer (PB) followed by fixation with 4% paraformaldehyde (PFA). Brains were removed and post-fixed for 1 hour in 4% PFA in PB at room temperature and were then dehydrated in 30% sucrose in 0.1 M PB. After the brains sank in the sucrose solution, they were frozen in 100% 2-methyl butane on dry-ice. Brains were sectioned on a sliding microtome at 40 µm and kept in cryoprotectant solution (25% glycerol, 25% propylene glycol in PBS) at -20°C until staining.

### Immunofluorescence

Immunofluorescence was performed as described in our previous studies (47, 48). Brain sections were washed in 0.1 M PBS 3 times for 5 minutes each before being placed into 100% methanol at -20°C for 10 minutes, immediately washed 3 times as before, and blocked in 4% normal goat serum with 0.4% Triton X-100 in 0.1M PBS for 1 hour. Sections were then incubated with primary antibody (anti-Phosphorylated STAT3, Cell Signaling, Danvers, MA, #9145, 1:200; anti-Tryptophan Hydroxylase (TPH), Millipore Darmstadt, Germany, AB1541, 1:750; anti-SERT, Neuromics, Edina, MN RA24330, 1:500; anti-5-HT, Neuromics, GT20079, 1:1,000) and incubated overnight at 4°C. The samples were then washed and incubated in secondary antibody (donkey anti-rabbit Alexafluor 488, Invitrogen, Carlsbad, CA A-21206 1:500; donkey anti-sheep Alexafluor 647, Invitrogen, A-21448 1:1,000; donkey anti-goat Alexafluor 555, Invitrogen A-21432 1:1000) for 4 hours at room temperature. Sections were washed, mounted on slides coated with 0.3% gelatin, and cover slipped with Vectashield antifade mounting medium (Vector Labs, Burlingame, CA; H-1000). All images were taken on a Leica SP8 confocal microscope.

### *In situ* hybridization

For RNAScope fluorescent in situ hybridization (FISH), animals were perfused with (PBS) and 4% PFA, and brains were processed as described above, but this time all the solutions were prepared in RNAse free conditions using autoclaved 0.1% DEPC treated water. Brains were sectioned on a cryostat at 20 µm and mounted on charged slides. Sections were then post fixed for 10 minutes in 4% PFA and dehydrated in increasing concentrations of ethanol. Then, the protocol for RNAScope Multiplex Fluorescent Reagent Kit v2 Assay was followed with the included reagents (Advanced Cell Diagnostics, Newark, CA) using probes against the leptin receptor mRNA (Cat# 415951) and Opal 650 dye kit (Akoya Biosciences, Marlborough, MA) to label LepR mRNA. Later, slides were washed in 0.05 M TBST then blocked in 10% normal horse serum and 1% bovine serum albumen (BSA) in 0.5 M TBS for 30 minutes. Primary antibody solution was then added (TPH: 1:500) and incubated overnight at 4°C. After 3 washes with TBS+Tween, Alexafluor 555 conjugated donkey anti-sheep secondary antibody (Invitrogen) was added (1:500) for 30 minutes. Slides were then washed and mounted with Prolong Gold antifade mounting medium (Thermofisher Scientific).

### *In vivo* microdialysis

Microdialysis was performed as described previously with minor modifications (49, 50). Rats were habituated to their own bowls in the BASi Raturn system for 20 hours over 4 consecutive days. Microdialysis sessions began the day after habituation ended, and the two sessions were separated by a 36-hour recovery. Each session was identical aside from the assigned treatment of each animal, vehicle or leptin. On the morning of the microdialysis session, rats were moved to the microdialysis room and fitted with probes from BASi (1 mm, MD-2202) into the ARC targeted cannula and perfused with artificial cerebral spinal fluid (150 mM NaCl, 3 mM KCl, 1.7 mM CaCl_2_·H_2_O, 0.183 mM MgCl_2_·6H_2_O, 5 mM D-Glucose) (ACSF) at a rate of 1 µL/minute. A brain infusion probe (1 mm, MD-2253) was also fitted into the DRN targeted cannula and connected to the treatment pump to not disturb the animal after collections had begun. The first three hours were discarded to allow recovery from the probe insertion. Dialysate samples were collected at 15-minute intervals into tubes containing 5 µL of antioxidant solution (0.5 mM acetic acid, 2.0 mM oxalic acid, 6.0 mM L-cysteine) and frozen at -80°C at the end of the collection. After the first 4 collections (baseline), rats were given DRN injections with either vehicle (sterile 0.1 M PBS) or 2 µg of leptin at 1 µL/min. Dialysates were collected for 4 hours post injection. Following the second day of microdialysis, animals were perfused, and the brains collected and sectioned as described above to verify probe placement in each rat.

### High performance liquid chromatography

Fifteen µL of microdialysate sample were run through an EICOMPAK PP-ODSII reversed phase analytical column, which isolates 5-HT from other biogenic compounds in interaction with a mobile phase consisting of 0.1 M Phosphate buffer pH 5.4 including 1.5% methanol, 500 mg/L decane-1-sulfonate sodium salt, and 50 mg/L 2Na-EDTA. 5-HT was measured at the graphite working electrochemical detector with an applied current of +400 mV and read using the EICOM HT-500 detector system. The concentrations of 5-HT in samples were interpolated against a three-point standard curve.

### Statistics

All statistical analyses were performed using GraphPad Prism 8 software. Simple comparisons between treatment and control groups were analyzed using a two-tailed unpaired t-test. Statistical analysis on the cumulative food intake as well as microdialysis data was done by two-way repeated measures analysis of variance (ANOVA) with time as a repeated measure, treatment as the independent variable. All analyses were followed by a Bonferroni post hoc test when appropriate, with α < 0.05 as the criterion for statistical significance. Values are reported as mean ± SEM.

## Acknowledgments

Supported by the National Science Foundation (IOS 1656626; CAG, JRF), NIH/NIGMS (P20 GM109091; CAG), NIH/NCATS (UL1TR00062; CAG), NIH (R01AG050518; JRF) and the Department of Veterans Affairs (IO1 BX001804; LPR).

## Notes

### Competing Interest Statement

The authors have declared no competing interest.

